# The chromatin insulator CTCF regulates HPV18 transcript splicing and differentiation-dependent late gene expression

**DOI:** 10.1101/2021.04.30.442078

**Authors:** Jack Ferguson, Karen Campos Leon, Ieisha Pentland, Joanne Stockton, Thomas Günther, Andrew Beggs, Sally Roberts, Boris Noyvert, Joanna L. Parish

**Author notes:** Equal Contribution (Jack Ferguson, Karen Campos Leon). Joint Senior Authors (Boris Noyvert, Joanna L. Parish). Corresponding Author: Joanna L. Parish.

## Abstract

The ubiquitous host protein, CCCTC-binding factor (CTCF), is an essential regulator of cellular transcription and functions to maintain epigenetic boundaries, stabilise chromatin loops and regulate splicing of alternative exons. We have previously demonstrated that CTCF binds to the E2 open reading frame (ORF) of human papillomavirus (HPV) 18 and functions to repress viral oncogene expression in undifferentiated keratinocytes by co-ordinating an epigenetically repressed chromatin loop within HPV episomes. Cellular differentiation, which is necessary for HPV life cycle completion disrupts CTCF-dependent chromatin looping of HPV18 episomes inducing enhanced activity of the HPV18 early promoter P_105_ and increased viral oncogene expression.

To further characterise CTCF function in HPV transcription control we utilised direct, long-read Nanopore RNA-sequencing which provides information on the structure and abundance of full-length transcripts. Nanopore analysis of primary human keratinocytes containing HPV18 episomes before and after synchronous differentiation allowed quantification of viral transcript species in these cultures, including the identification of low abundance novel transcripts. Comparison of transcripts produced in wild type HPV18 genome-containing cells to those identified in CTCF-binding deficient genome-containing cells (HPV18-ΔCTCF) identifies CTCF as a key regulator of differentiation-dependent late promoter activation, required for efficient E1^E4 and L1 protein expression. Furthermore, our data show that CTCF binding at the E2 ORF of HPV18 promotes usage of the downstream weak splice donor (SD) sites SD3165 and SD3284, to the dominant E4 splice acceptor site at nucleotide 3434. These findings demonstrate importance of CTCF-dependent transcription regulation at multiple stages of the HPV life cycle.

**IMPORTANCE:** Oncogenic human papillomavirus (HPV) infection is the cause of a subset of epithelial cancers of the uterine cervix, other anogenital areas and the oropharynx. HPV infection is established in the basal cells of epithelia where a restricted programme of viral gene expression is required for replication and maintenance of the viral episome. Completion of the HPV life cycle is dependent on the maturation (differentiation) of infected cells which induces enhanced viral gene expression and induction of capsid production. We previously reported that the host cell transcriptional regulator, CTCF, is hijacked by HPV to control viral gene expression. In this study, we use long-read mRNA sequencing to quantitatively map the variety and abundance of HPV transcripts produced in early and late stages of the HPV life cycle and to dissect the function of CTCF in controlling HPV gene expression and transcript processing.

## INTRODUCTION

Human papillomaviruses (HPVs) are a family of small, double-stranded DNA viruses that infect cutaneous and mucosal epithelia. Most HPV types cause benign epithelial hyperproliferation, which is usually resolved by host immune activation. However, persistent infection with a subset of HPV types (e.g., HPV16 and 18) are the cause of epithelial tumours including cervical and other anogenital cancers, and carcinoma of the oropharyngeal tract (1).

The viral episome is maintained and replicated in the cell nucleus as an extrachromosomal, chromatinised episome which allows the epigenetic regulation of viral transcription in an equivalent manner to host genes (2). The regulation of HPV gene expression in differentiating epithelia is tightly regulated and is a key strategy in the maintenance of persistent infection. Several distinct transcriptional start sites (TSSs) have been identified including the major early and late promoters, the E8 promoter (P_E8_) and less well-defined TSSs around nucleotide 520 (P_520_) and 3000 (P_3000_). The relative activity of these promoters is dependent on the differentiation status of the host keratinocyte (3–5). Establishment of HPV infection occurs in the undifferentiated basal keratinocytes of epithelia where viral genome copy number and transcription are maintained at low levels, presumably to prevent host immune activation. We and others have shown that the viral episome is maintained in an epigenetically repressed state in undifferentiated keratinocytes, characterised by low abundance of trimethylation of lysine 4 (H3K4Me3) and enrichment of trimethylation of lysine 27 (H3K27Me3) on histone H3, which attenuates viral gene expression (5, 6). The host cell chromatin-organising and transcriptional insulation factor, CCCTC-binding factor (CTCF) is important in the maintenance of the epigenetic repression of the HPV genome through the stabilisation of a chromatin loop. CTCF binds to a conserved site in the E2 open reading frame (ORF) of HPV18 approximately 3,000 base pairs downstream of the viral transcriptional enhancer situated in the long control region (LCR) (7). Although the major CTCF binding site and the viral enhancer are physically separated, we demonstrated that abrogation of CTCF binding resulted in inappropriate epigenetic activation of the HPV18 enhancer and early promoter (termed P_105_ in HPV18) and increased expression of the viral oncoproteins E6 and E7 (E6/E7) (6, 7). CTCF physically associates with the transcriptional repressor Ying Yang 1 (YY1) (8) and we subsequently showed that CTCF-dependent epigenetic repression of the HPV18 episome was through interaction with YY1 bound at the viral LCR, such that CTCF and YY1 co-operate to stabilise an epigenetically repressed chromatin loop within the early gene region (6). While the association of CTCF with the HPV18 episome is not significantly altered by keratinocyte differentiation, YY1 protein expression and binding to the HPV18 genome is dramatically reduced in differentiated keratinocytes leading to loss of CTCF-YY1 dependent chromatin loop stabilisation (6). This differentiation-dependent topological change in the HPV episome is coincident with epigenetic activation of the P_105_ promoter and increased expression of the HPV E6/E7 oncoproteins.

Activation of the major late promoter (termed P_811_ in HPV18) in part occurs through epigenetic derepression of the HPV episome upon keratinocyte differentiation (5, 6, 9) and reviewed in (10). This restricts expression of the viral capsid proteins L1 and L2 to the upper compartment of infected epithelia, limiting their potential for host immune activation (4, 11, 12). The late promoter also regulates expression of viral intermediate genes including E1, E2, E1^E4 and E5, which are important for viral genome amplification in the upper layers of the infected epithelia (13, 14). The mechanisms underlying the differentiation-dependent epigenetic activation of late promoter activity are not clear, but it has been shown that the viral enhancer in the LCR is required for late promoter activation (15) and that differentiation-dependent enhancement of transcription elongation may play a key role in late promoter activation (16).

Further enhancing the complexity of HPV gene expression regulation, the polycistronic HPV mRNA is subject to extensive post-transcriptional splicing, which gives rise to an array of transcripts that each encode a distinct subset of full length, and/or fusion proteins. While studies have mapped the HPV18 transcriptome (17, 18), the quantification of HPV promoter activity and the abundance of each mature transcript has not been reported. Cellular splicing factors are utilised and manipulated by the virus to co-ordinate differentiation dependent viral transcript splicing, including the serine-arginine rich (SR) proteins and heterogeneous ribonucleoproteins (hnRNPs) (19, 20). In addition to its functions in chromatin looping and epigenetic isolation, CTCF can play an important role in regulating alternative gene splicing, most likely through multiple mechanisms. In the host cell *CD45* locus, CTCF binding within exon 5 promotes inclusion of upstream exons by creating a “roadblock” to pause RNA polymerase II progression, allowing more efficient recognition of weak exons by the splicing machinery (21). It has also been shown that DNA methylation-dependent binding of CTCF within normally weak exons promotes inclusion during co-transcriptional splicing (22). To support these findings, a significant enrichment of CTCF binding sites in close proximity to alternatively spliced exons has been reported (23). However, CTCF binding at distant sites can also influence alternative exon usage through the stabilisation of intragenic chromatin loops (24). Our early analysis of CTCF-dependent control of HPV18 transcript splicing indicated an important role for this factor in maintaining the complexity of slicing events (7) but the global effect of CTCF on HPV18 transcript processing was not analysed.

Next generation sequencing (NGS) has revolutionised virology research by providing nucleotide resolution data on existing and emerging pathogens, prevalence, and evolution. However, conventional Illumina-based RNA sequencing (RNA-Seq) methods are limited in that information on the structure of full-length transcripts, including alternative splicing is sacrificed to preserve accuracy and read depth (25). Direct, long-read Nanopore sequencing overcomes this limitation by providing quantitative data on the abundance of individual mRNA isoforms (26).

In this study, we use Nanopore sequencing to quantify the spectrum of HPV18 transcripts in HPV18 episome-containing primary human keratinocytes and to map differentiation-induced changes in promoter usage, splicing and transcript abundance. Furthermore, we characterise the global effect of CTCF binding to the HPV18 genome on transcript splicing and early and late promoter activity.

## METHODS

### Ethical approval

The collection of neonatal foreskin tissue for the isolation of primary human foreskin keratinocytes (HFKs) for investigation of HPV biology was approved by Southampton and South West Hampshire Research Ethics Committee A (REC reference number 06/Q1702/45). Written consent was obtained from the parent/guardian. The study was approved by the University of Birmingham Ethical Review Committee (ERN 16-0540).

### Cell culture, methylcellulose differentiation and organotypic raft culture

Normal primary HFKs from neonatal foreskin epithelia were transfected with recircularised HPV18 wild type (WT) or -ΔCTCF genomes and maintained on irradiated J2-3T3 fibroblasts in complete E medium (27) as previously described (7). For methylcellulose-induced keratinocyte differentiation, 3 x10^6^ HPV18-WT or - ΔCTCF genome containing keratinocytes were suspended in E-media supplemented with 10 % FBS and 1.5 % methylcellulose and incubated at 37 °C, 5 % CO_2_ for 48 hrs. Cells were then harvested by centrifugation at 250 x g followed by washing with ice-cold PBS. Cells were then either suspended in medium containing 1 % formaldehyde to cross-link for chromatin immunoprecipitation (ChIP) as described below, or RNA and protein was extracted from cell pellets as previously described (7).

Organotypic raft cultures were prepared as previously described (7). Rafts were cultured for 14 days in E medium without epidermal growth factor to allow cellular stratification. Raft cultures were fixed in 3.7 % formaldehyde and paraffin embedded and sectioned by Propath Ltd (Hereford, United Kingdom).

### Antibodies

Anti-CTCF (61311) and anti-H4Ac (39925) antibodies was purchased from Active Motif, HPV18 L1 (ab69) antibody purchased from Abcam and anti-β-actin (AC-74) purchased from Sigma. E1^E4 antisera were produced as previously described (28). HRP-conjugated anti-mouse and anti-rabbit antibodies (Jackson Laboratories) were used for Western blotting and Alexa-488 and −594 conjugated anti-rabbit/mouse antibodies (Invitrogen) were used for immunofluorescence staining.

### Chromatin immunoprecipitation-qPCR (ChIP-qPCR)

ChIP-qPCR assays were performed using the ChIP-IT Express Kit (Active Motif) as per the manufacturer’s protocol. Briefly, cells were fixed in 1 % formaldehyde for 5 mins at room temperature with gentle rocking, quenched in 0.25 M glycine and washed with ice-cold PBS. Nuclei were released using a Dounce homogeniser. Chromatin shearing was carried out by sonication at 25 % amplitude for 30 secs on/30 secs off for a total time of 15 mins using a Sonics Vibracell sonicator fitted with a microprobe. ChIP efficiency was assessed by qPCR using SensiMix SYBR master mix using a Stratagene Mx3005P (Agilent Technologies, Santa Clara, CA, USA). Primer sequences for ChIP experiments are shown in Table 1. Cycle threshold (C_T_) values were used to calculate fold enrichment compared to a negative control FLAG antibody with the following formula:

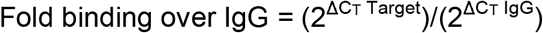

**Table 1:**
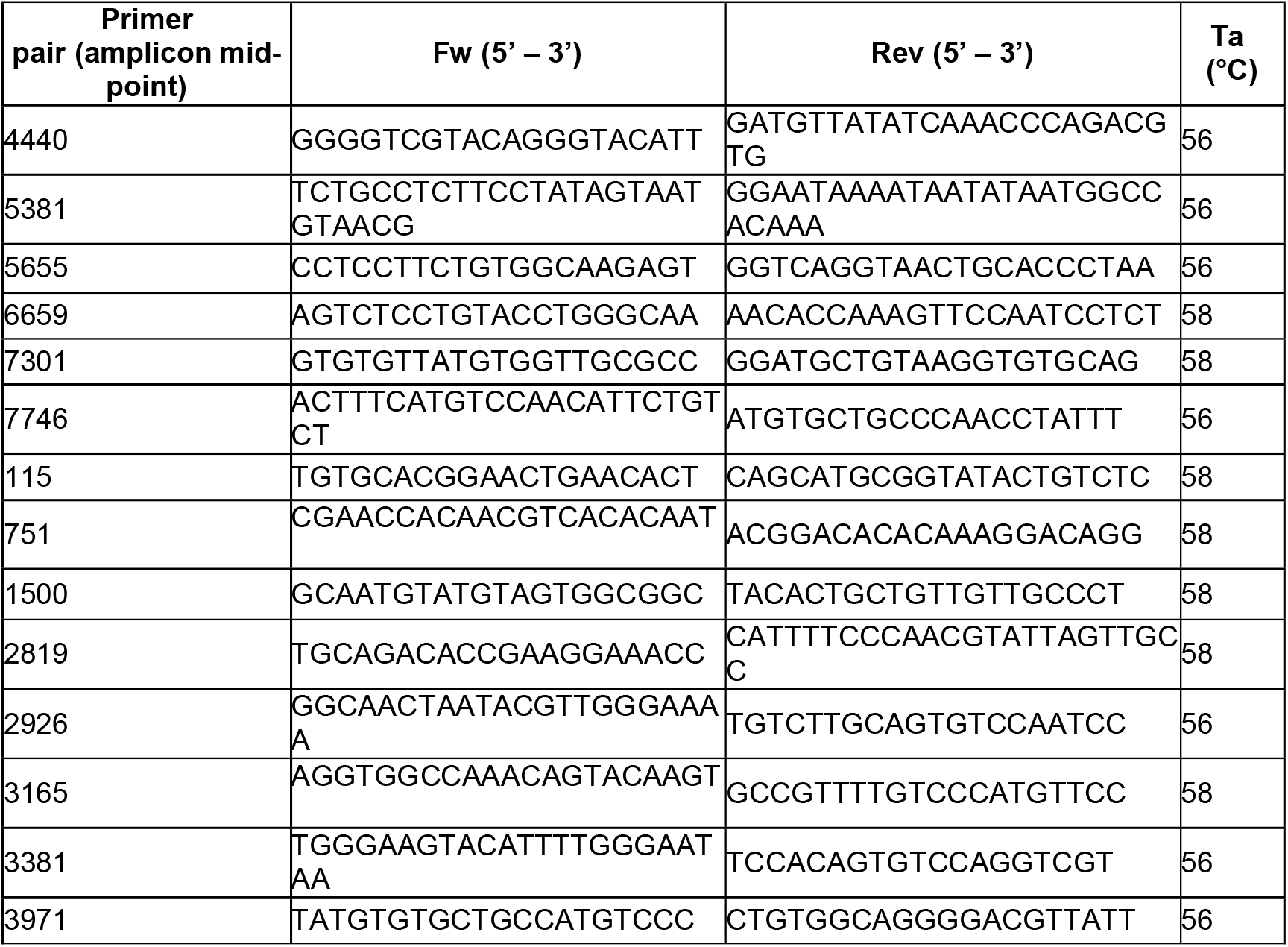
Primer sequences used for ChIP-qPCR experiments. Ta, annealing temperature

Where ΔC_T_ target = Input C_T_ – Target C_T_ and ΔC_T_ IgG = Input C_T_ – IgG C_T_. Each independent experiment was performed in technical triplicate and data shown are the mean and standard deviation of three independent repetitions.

### ChIP-Seq

ChIP and respective input samples were used for generation of ChIP-Seq libraries as described (29). Briefly, 2-10 ng DNA was used in conjunction with the NEXTflex Illumina ChIP-Seq library prep kit (Cat# 5143-02) as per the manufacturer’s protocol. Samples were sequenced on a HiSeq 2500 system (Illumina) using single read (1×50) flow cells. Sequencing data was aligned to the HPV18 genome (accession number: AY262282.1) using Bowtie (30) with standard settings and the -m1 option set to exclude multi mapping reads (31).

### RNA sequencing and data analysis

For RNA-Seq, libraries were prepared using Tru-Seq Stranded mRNA Library Prep kit for NeoPrep (Illumina, San Diego, CA, USA) using 100ng total RNA input according to manufacturer’s instructions. Libraries were pooled and run as 75-cycle–pair end reads on a NextSeq 550 (Illumina) using a high-output flow cell. Sequencing reads were aligned to human (GRCh37) and HPV18 (AY262282.1) genomes with STAR aligner (v2.5.2b) (32). The computations were performed on the CaStLeS infrastructure (33) at the University of Birmingham. Sashimi plots were generated in Integrative Genomics Viewer (IGV), Broad Institute ((http://software.broadinstitute.org/software/igv/).

### Nanopore direct RNA sequencing and data analysis

8×10^7^ cells from undifferentiated or methylcellulose differentiated keratinocytes containing HPV18 (WT or ΔCTCF) samples for RNA extraction using the RNeasy Plus Mini Kit (Qiagen) according to the manufacturer’s instructions and DNaseI treated (Promega). 500 ng of polyA+ RNA was used in conjunction with the direct RNA sequencing kit (Oxford Nanopore technologies, Oxford, UK [SQK-RNA002]). All protocol steps are as described in (34). The reads were aligned to the human (GRCh37) and HPV18 (AY262282.1) genomes using minimap2 (35) with options “-ax splice -uf -k14” for nanopore direct RNA mapping. The splicing coordinates were extracted from the bam files using custom scripts.

### Cell lysis and western blotting

Cells were lysed with urea lysis buffer (ULB; 8 M urea, 100 mM Tris-HCl, pH 7.4, 14 mM ß-mercaptoethanol, protease inhibitors) and protein concentration determined. Protein extracts from organotypic raft cultures were harvested using ULB and homogenised using a Dounce homogeniser contained with a category II biological safety cabinet. Lysates were incubated on ice for 20 mins before centrifugation at 16,000 x g for 20 mins at 4 °C. Supernatant was transferred to a fresh tube and protein concentration assessed by Bradford Assay. For Western blotting, equal quantities of protein lysates were separated by SDS-PAGE and western blotting was carried out by conventional methods. Chemiluminescent detection was carried out using a Fusion FX Pro and densitometry performed with Fusion FX software.

### Immunofluorescence

Immunofluorescence was carried out on paraffin embedded organotypic raft culture sections using the agitated low temperature epitope retrieval (ALTER) method as previously described (36). Briefly, slides were sequentially immersed in Histoclear (Scientific Laboratory Supplies) and 100 % IMS and incubated at 65 °C in 1 mM EDTA (pH 8.0), 0.1 % Tween 20 overnight with agitation. Slides were then blocked in PBS containing 20 % heat-inactivated normal goat serum and 0.1 % BSA (Merck). Primary antibodies were diluted in block solution and incubated overnight at 4 °C followed by 3x PBS washes. Fluorophore-conjugated secondary antibodies were diluted in block buffer and added to slides which were incubated at 37 °C for 1 hour. Slides were subsequently washed 4x 10 mins in PBS with Hoechst 33342 solution (10 μg/ml) added to the final PBS wash. Slides were mounted in Fluoroshield (Sigma-Aldrich) and visualised using a Nikon inverted Epifluorescent microscope fitted with a 40x oil objective. Images were captured using a Leica DC200 camera and software.

## Results

We have previously characterised a CTCF binding site within the E2 open reading frame (ORF) of HPV18 which is strongly bound by CTCF in a primary HFK model of the HPV18 life cycle (6, 7). Although the E2-CTCF binding site was the most CTCF enriched region of the HPV18 genome in our ChIP-qPCR analysis, there did appear to be other regions of the viral genome that were bound at a lower level by CTCF. In addition, CTCF binding sites have been predicted in the late gene region of HPV18 and other high-risk HPV types and binding has been demonstrated in HPV31 episomes (7, 37). To analyse CTCF binding to the HPV18 genome with greater sensitivity, we opted to map CTCF binding peaks using ChIP-sequencing (ChIP-Seq). Anti-CTCF immunoprecipitated chromatin harvested from HFKs harbouring HPV18 episomes was subject to Illumina next generation sequencing. Reads were aligned to the HPV18 genome revealing robust enrichment of CTCF in the E2 ORF with maximal binding between nucleotides 2960-3020, corresponding to the previously identified E2-CTCF binding site (**Fig.1A**). No other distinct CTCF peaks were observed in the HPV18 genome. In addition, ChIP-Seq analysis of CTCF enrichment in mutant HPV18 genomes in which the E2-CTCF binding site was mutated to prevent CTCF binding (HPV18-ΔCTCF), revealed a complete loss of CTCF binding to the E2-ORF with no evidence of enhanced binding at secondary sites (**Fig.1A**), confirming our previous ChIP-qPCR analysis of this mutant virus.

**Figure 1:**
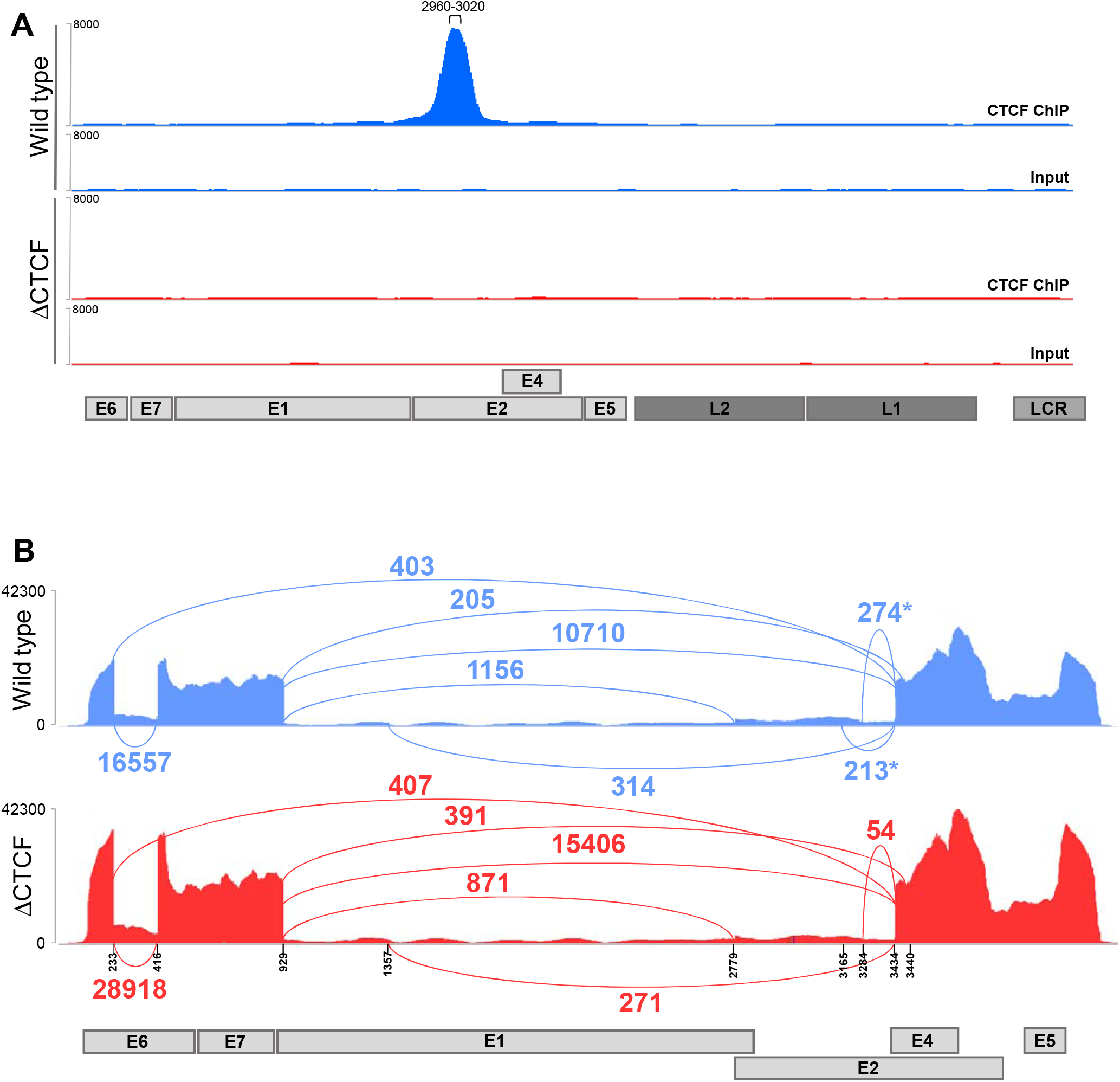
Abrogation of CTCF recruitment to the HPV18 E2 ORF alters early transcript splicing. (A) Enrichment of CTCF in the HPV18 genome was assessed by ChIP-Seq in either WT (blue) or ΔCTCF-HPV18 (red) genome-containing keratinocytes. Next generation sequencing data were IGV. The position of HPV18 ORFs and LCR are indicated below the alignment profiles. (B) Exon-exon junctions in Illumina-based RNA-Seq data sets of either WT (blue) or ΔCTCF-HPV18 (red) genome-containing keratinocytes were identified and quantified in IGV and represented in Sashimi plots. The co-ordinates of splice donor and acceptor sites and annotated ORFs are indicated. The number of reads at each exon-exon junction is indicated. *denotes splicing event identified in WT HPV18 but reduced or lost in ΔCTCF-HPV18 genome containing cells.

Abrogation of CTCF binding at the HPV18 E2 ORF resulted in increased transcriptional activity of the HPV18 early promoter (P_102_) and a concomitant increase in E6/E7 protein expression (6, 7). These studies also revealed alterations in the splicing of early transcripts, indicated by a significant reduction in the abundance of transcripts spliced at 233^3434 upon amplification by semi-quantitative RT-PCR (7). To confirm these findings and to further characterise CTCF-dependent regulation of HPV18 transcript splicing, we utilised high-depth Illumina RNA-Seq data in HPV18-wild type and -ΔCTCF transfected primary HFKs to quantify individual splicing events (**Fig.1B**). While there were a similar number of splicing events at 233^3434 in the wild type and mutant HPV18 genome-containing cells (403 and 407 events, respectively), splicing at 233^416 was increased in HPV18-ΔCTCF genome containing cells in comparison to wild type (28,918 events compared to 16,557 events respectively, Fisher’s test p-value <0.00001), which could account for the observed relative reduction in amplification of transcripts spliced at 233^3434 by qRT-PCR (7). Interestingly, we also noted a reduction in splicing at 3284^3434, previously proposed to encode a truncated form of the E2 protein, E2C (18), and a complete loss of splicing at 3165^3434 in HPV18-ΔCTCF genome containing cells compared to wild type HPV18. Found at relatively low abundance, splicing at 3165^3434 has been previously described and predicted to encode a novel E2^E4 fusion protein termed E2^E4L (38). Similarly, splicing at 2853^3434 has been proposed to encode a shorter form of E2^E4 fusion protein, E2^E4S (38), however, this splice was not detected in our Illumina RNA-Seq data. Splicing from the 929 SD site to the weak acceptor site at 3440 was observed in HPV18-ΔCTCF genome containing cells (391 events) but not in wild type HPV18. These findings suggest that CTCF may play a role in controlling acceptor site usage downstream of the E2-CTCF binding site.

While individual splicing events can be quantified using conventional short-read RNA sequencing methods, the evaluation of the structure of individual transcripts and the multiple splicing events that may occur within a single transcript is not possible. To fully characterise and, for the first time, quantify the relative abundance of individual HPV18 transcripts in primary HFKs, purified and polyA+ enriched RNA was analysed by direct long-read MinION sequencing. Cells were either grown in monolayer culture on feeder cells (undifferentiated) or embedded in semi-solid methylcellulose containing medium for 48 hours, to induce synchronous differentiation, and the spectrum of transcripts quantified as reads per million (RPM) for each sample. To confirm that morphological keratinocyte differentiation was induced by suspension in methylcellulose, differentiation-dependent changes to cellular markers of keratinocyte differentiation were analysed. A notable increase in involucrin (IVL) expression was observed (**Fig.2A;** Fisher’s test p-value < 0.00001). In addition, an alteration in transcript splicing of the keratinocyte-specific extracellular matrix protein, Ecm1, upon keratinocyte differentiation has been reported (39). Undifferentiated keratinocytes express full length Ecm1 transcript 2 but expression of a shorter, alternatively spliced transcript (transcript 3) is induced upon keratinocyte differentiation. Analysis of Ecm1 transcripts in our MinION sequencing data demonstrated the appearance of Ecm1 transcript 3 which lacks exon 7 in methylcellulose differentiated keratinocytes only (**Fig.2B**).

**Figure 2:**
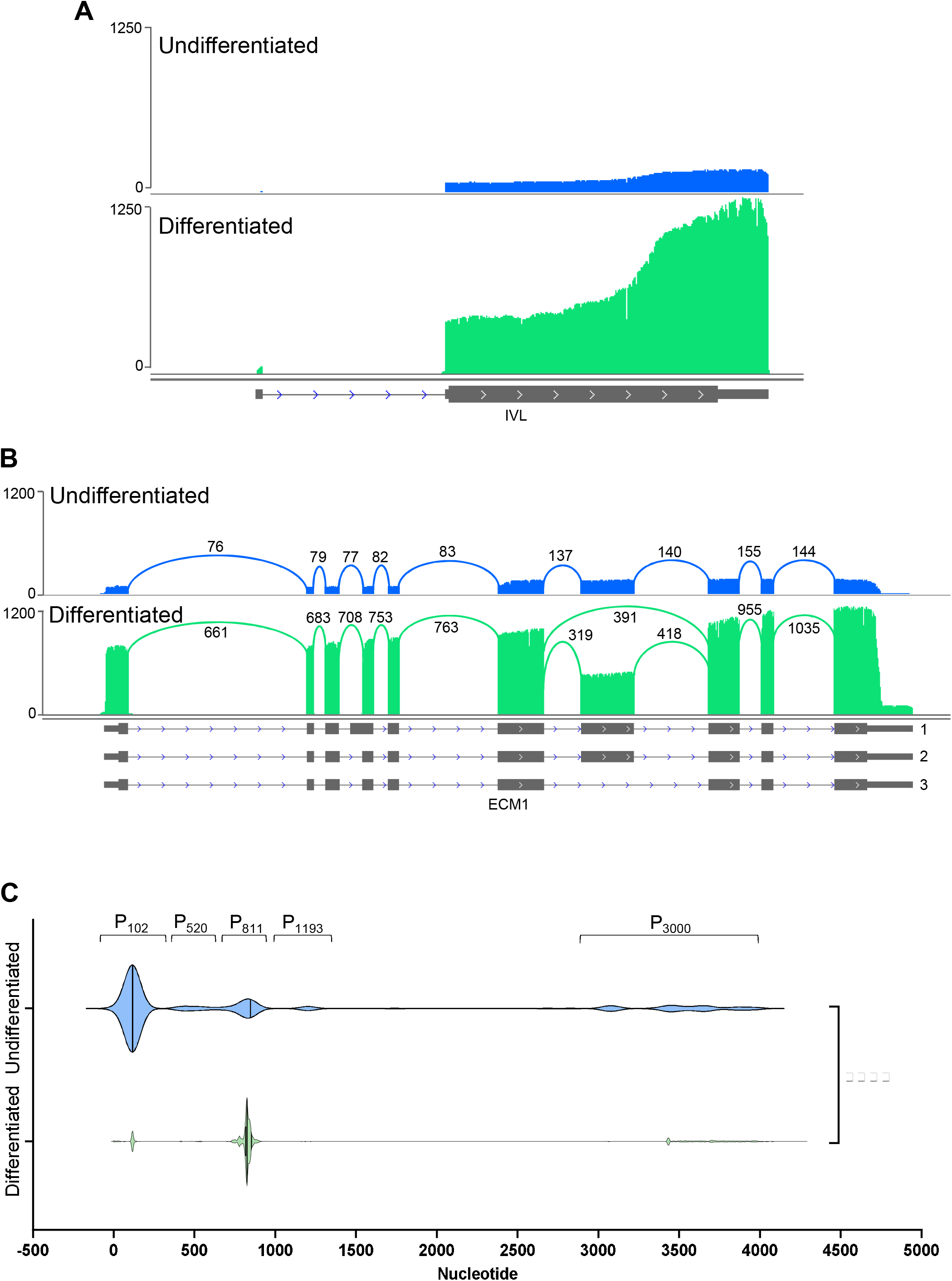
Analysis of differentiation-dependent host cell gene expression and HPV transcriptional start site usage. HPV18-HFK were synchronously differentiated in methylcellulose for 48 hrs (green). The host and viral transcriptomes were analysed by long read RNA-Seq and compared to undifferentiated HPV18-HFK (blue). (A) Enhanced involucrin (IVL) expression following keratinocyte differentiation and (B) enhanced ECM1 expression combined with differentiation-induced exon 7 skipping. (C) Clustered HPV18 TSS usage in undifferentiated and differentiated keratinocytes showing differentiation-dependent alteration of the major early (P_102_) and major late (P_811_) promoter usage. ****p < 0.0001 (Fisher’s test).

Virus host fusion transcripts were identified at very low abundance (<2% of total HPV reads), indicative of low-level viral integration. Nonetheless, these fusion transcripts were removed from our data set prior to analysis to include only those transcripts derived from HPV episomes. Data were then normalised to the total number of reads in each sample to calculate reads per million (RPM) of each viral transcript species. In agreement with previous reports (17, 18), five clear groupings of transcriptional start sites were identified in undifferentiated HPV18 genome containing cells, which originated between nucleotides 1-350 (P_102_), 351-700 (P_520_), 701-900 (P_811_), 1000-1400 (P_1193_) and 2800-4000 (P_3000_) (**Fig.2C**), which were used to define transcript species in subsequent quantifications. Keratinocyte differentiation resulted in a significant change in TSS usage characterised by activation of the P_811_ major late promoter (**Fig.2C**). In undifferentiated HPV18 wild type genome-containing cells, the most abundant transcript was initiated at the P_102_ promoter and spliced at 233^416-929^3434 (transcript 3; **Fig.3**). This transcript has the potential to encode E6*I, E7, E1^E4 and E5. Several novel transcripts were identified at low abundance in the undifferentiated wild type HPV18 cells including transcripts 13 and 25, which encode E6*I and E7 respectively along with E2 and E5. Interestingly, splicing at both 3165^3434 and 3284^3434 was observed in undifferentiated cells (transcripts 11 and 12; **Fig.3**), however, these transcripts originated from the P_3000_ promoter and therefore lack the E2 start codon at nt2816 and more likely encode E5 in the basal keratinocytes rather than E2^E4 fusion proteins as previously suggested (38). A single, previously described transcript, spliced at 929^2779-3165^3434 was identified in differentiated WT HPV18 genome containing cells that originated at the P_105_ promoter, which contains the E2 start codon and could therefore encode E2^E4L (transcript 6; **Fig.3**)(38). We also identified very low abundance of transcripts spliced at 929^2779-3284^3434, which is in frame with the E4 ORF and we predict would encode a novel E2^E4 fusion protein, denoted E2^E4XL but the presence of this theoretical protein in HPV18 infected cells is yet to be confirmed (transcript 5; **Fig.3**).

**Figure 3:**
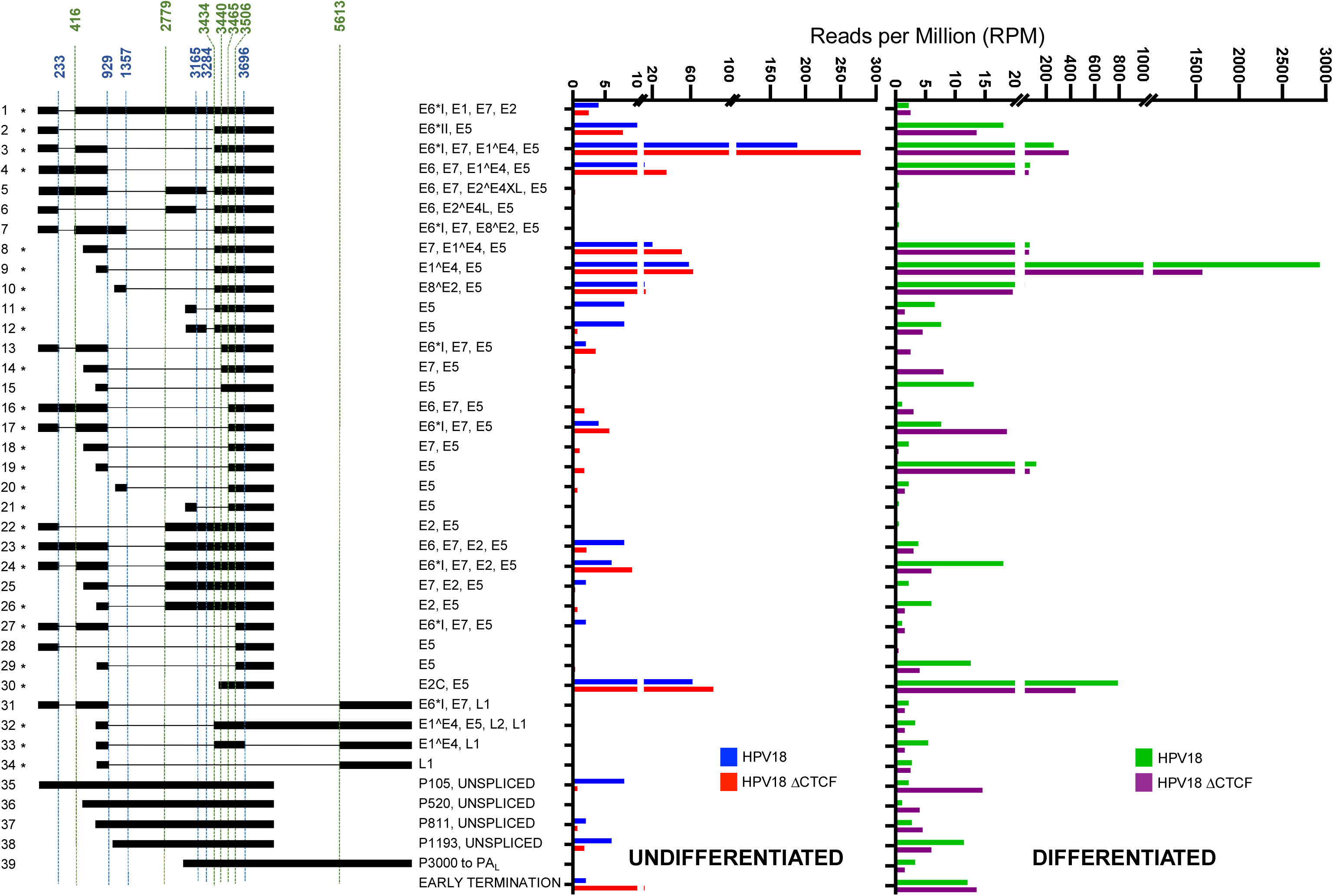
Quantitative analysis of the HPV18 transcriptome in undifferentiated and differentiated keratinocytes and alterations induced by abrogation of CTCF binding. Alignment of direct RNA sequencing data to the HPV18 genome facilitated the characterisation of all HPV-specific transcripts. Transcripts were only included in the data set if they were represented by two or more reads. The relative abundance of each transcript type was calculated in reads per million (RPM) of the total reads in each sample. Relative abundance (RPM) of each transcript is shown for WT (blue) and ΔCTCF-HPV18 (red) genome-containing cells in undifferentiated keratinocytes (left) and for WT (green) and ΔCTCF-HPV18 (purple) in differentiated keratinocytes (right). The identified splice donor (blue) and acceptor (green) sites are indicated above the transcript map and HPV18 ORFs encoded by each transcript are shown. *denotes transcripts that have previously been identified (17, 18).

Comparison of viral transcripts in WT and HPV18-ΔCTCF genome-containing cells revealed a significant increase in abundance of the major early transcript originating from the P_105_ promoter and spliced at 233^416-929^3434, which encodes E6*I, E7, E1^E4 and E5 (transcript 3; **Fig.3**, Fisher’s test p-value < 0.00001). A more modest increase in the second most abundant transcript in undifferentiated cells, originating from the P_105_ promoter and spliced at 929^3434 was also observed, which has the potential to encode full length E6 as well as E7, E1^E4 and E5 (transcript 4; **Fig.3**, Fisher’s test, non-significant). The increased abundance of these major early viral transcripts corroborates the previously observed increase in E6 and E7 protein expression when CTCF binding site is ablated (6, 7).

Notably, splicing at both 3165^3434 and 3284^3434 (transcripts 11 and 12; **Fig.3**) was significantly reduced in HPV18-ΔCTCF genome containing cells compared to WT (Fisher’s test p-value < 0.00001 and 0.01, repectively) corroborating our finding in Illumina RNA-Seq datasets that CTCF may function to enhance the activity of downstream weak SD sites in the HPV18 genome. Transcripts spliced at 929^3440 (transcripts 13, 14 and 15) were also detected at low abundance.

Transcripts that originate from the P_811_ late promoter were abundantly expressed in undifferentiated cells; transcripts originating from this promoter and spliced at 929^3434 to encode E1^E4 and E5 proteins (transcript 9; **Fig.3**) were the second most abundant transcript in undifferentiated cells. As expected, the abundance of this transcript was dramatically increased around 50-fold (Fisher’s test p-value < 0.00001) upon differentiation of the WT HPV18 cells in methylcellulose. However, while differentiation of HPV18-ΔCTCF genome-containing cells similarly resulted in an increase in abundance of this major E1^E4 encoding transcript, the overall abundance of this transcript was reduced by around 50 % compared to WT. It is also interesting to note that transcripts encoding the L1/L2 capsid proteins (transcripts 31-34; **Fig.3**) were induced upon cellular differentiation in WT genome-containing cells, albeit at a low level, but these transcripts were all lower in abundance in HPV18-ΔCTCF cells. These data suggest that recruitment of CTCF to the HPV18 genome at the E2-ORF may be important for differentiation-dependent activation of the viral late promoter.

The major transcriptional promoters in the HPV18 genome have been previously mapped using 5’ RACE (17). Although transcript sequencing by Nanopore does not provide nucleotide resolution accuracy in mapping transcription start sites (40), the clustering of the 5’ end of viral transcripts was clearly enriched at the previously annotated transcriptional start sites (**Fig.2C**). Therefore, to characterise the differential activity of the major viral promoters in HPV18 WT and -ΔCTCF cells, the 5’ end of each viral read in our Nanopore datasets was mapped and quantified. The 5’ end of most transcripts (>95 %) mapped near four previously described promoters; P_102_, P_520_, P_811_ and P_3000_ (**Fig.4**). Interestingly, the 5’ end of transcripts that originated from both the P_102_ and P_811_ promoters clustered as a sharp peak at the previously annotated transcriptional start site whereas the 5’ end of transcripts originating from either the P_520_ or P_3000_ promoter regions were more broadly distributed (**Fig.4A-D**). As expected, the P_102_ promoter was the most active promoter in HPV18 WT genome-containing undifferentiated cells with very few transcripts originating from the P_811_ late promoter. Differentiation of these cells resulted in a dramatic increase in transcripts originating from the P_811_ promoter (Fisher’s test p-value < 0.00001), coincident with a slight increase in P_102_ activity (Fisher’s test p-value < 0.00001) (**Fig.4A and C**). Transcripts originating from the P_102_ promoter were ~30 % more abundant in HPV18-ΔCTCF genome containing cells than WT, which was further activated upon cellular differentiation confirming enhanced activity of the early promoter in the absence of CTCF recruitment. Interestingly, the activity of the P_811_ late promoter was notably lower in differentiated HPV18-ΔCTCF genome containing cells compared to WT (Fisher’s test p-value < 0.00001), providing evidence that the activity of the late promoter in differentiated cells is attenuated when CTCF recruitment is abrogated. The P_520_ promoter had reduced activity compared to the P_102_ and P_811_ promoters, but interestingly this promoter was more active in differentiated HPV18 WT genome-containing cells than undifferentiated (Fisher’s test p-value 1.8E-4). In HPV18-ΔCTCF cells, the P_520_ promoter was more active than WT in undifferentiated cells (Fisher’s test p-value 0.016) but was not further activated by cellular differentiation (**Fig.4B**). Very few transcripts originated from P_3000_ in undifferentiated cells, however this promoter was strongly activated following cellular differentiation in HPV18 WT genome containing cells. As was observed at P_811_, differentiation-dependent activation of P_3000_ was reduced in HPV18-ΔCTCF genome containing cells compared to WT. The P_E8_ promoter (P_1193_) was only weakly active with less than 3 % of transcripts originating at this promoter in undifferentiated cells. Furthermore, the activity of P_E8_ was not affected by keratinocyte differentiation or mutation of the E2-CTCF binding site (data not shown). Analysis of TSS usage in the bulk population of viral transcripts revealed that while there was a greater proportion of transcripts which initiated from the P_102_ early promoter in HPV18-ΔCTCF episomes than WT (indicated by tighter density grouping), this did not reach significance (p 0.16) (**Fig.5A**). In contrast, highly significant differences were observed between TSS usage in HPV18-ΔCTCF episomes compared to WT following keratinocyte differentiation (p < 1E-16). While in WT HPV18 cells, the TSS usage density was highly enriched at the P_811_ promoter, transcripts in ΔCTCF-HPV18 genome-containing cells were less abundant at the P_811_ promoter, and the P_102_ promoter was proportionately more active than in WT-HPV18 episomes (**Fig.5B**). These analyses demonstrate that differentiation-dependent stimulation of P_811_ major late promoter activity is facilitated by recruitment of CTCF to the E2 ORF.

**Figure 4:**
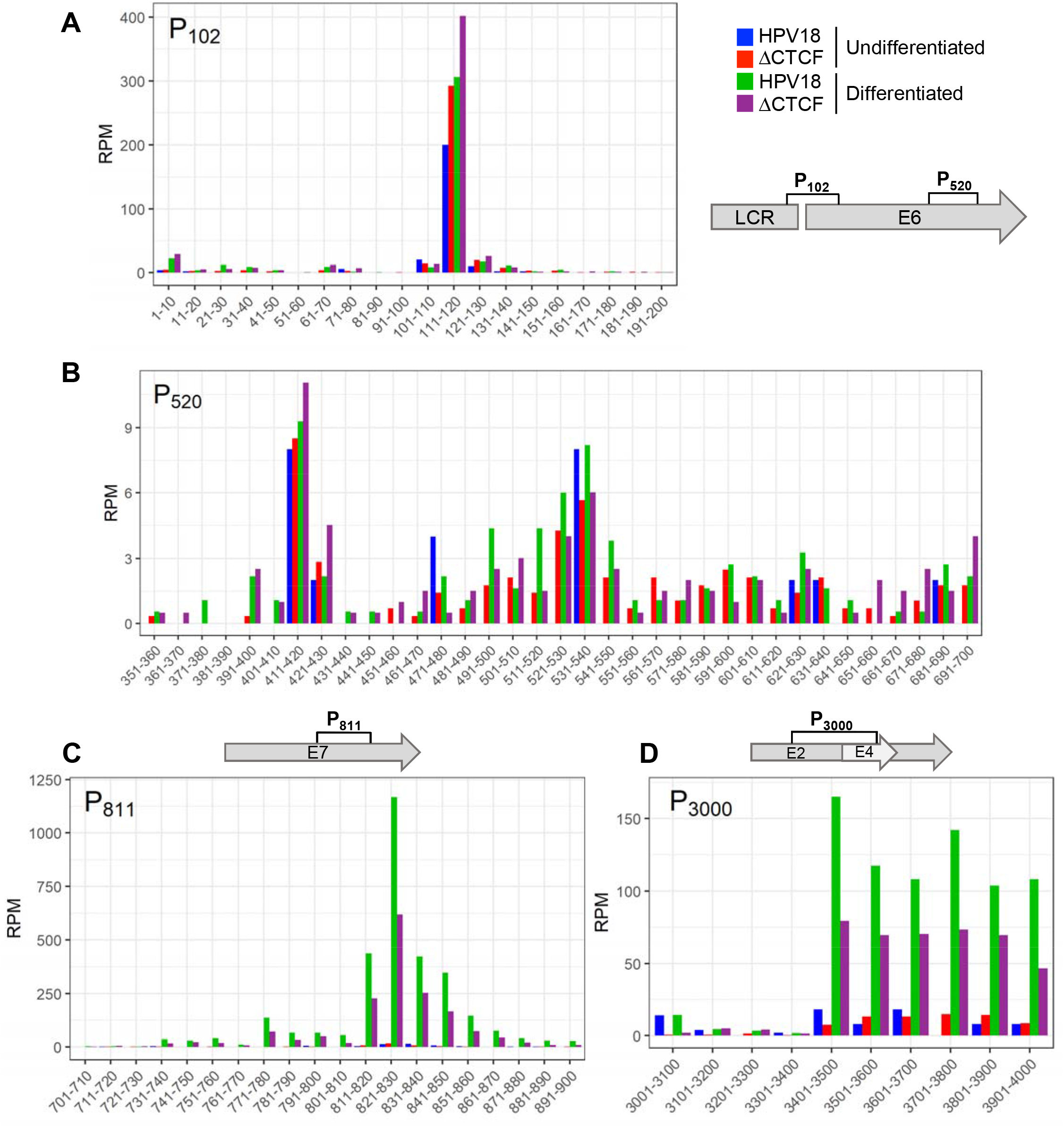
Quantitative analysis of transcription start site usage in undifferentiated and differentiated keratinocytes and CTCF-dependent regulation of promoter activity. The 5’ end of each HPV18 transcript was identified in Nanopore RNA sequencing data sets and relative abundance calculated as reads per million (RPM). Total counts at each nucleotide position were binned into 10 (A, B, C) or 100 (D) nucleotide regions in the data shown. Transcripts originating around the P_105_ (A), P_520_ (B), P_811_ (C) and P_3000_ (D) promoters were identified in wild type and ΔCTCF-HPV18 cells in undifferentiated (blue and red, respectively) and methylcellulose differentiated (green and purple, respectively) cultures. Relevant HPV18 genome features are shown alongside each panel.

**Figure 5:**
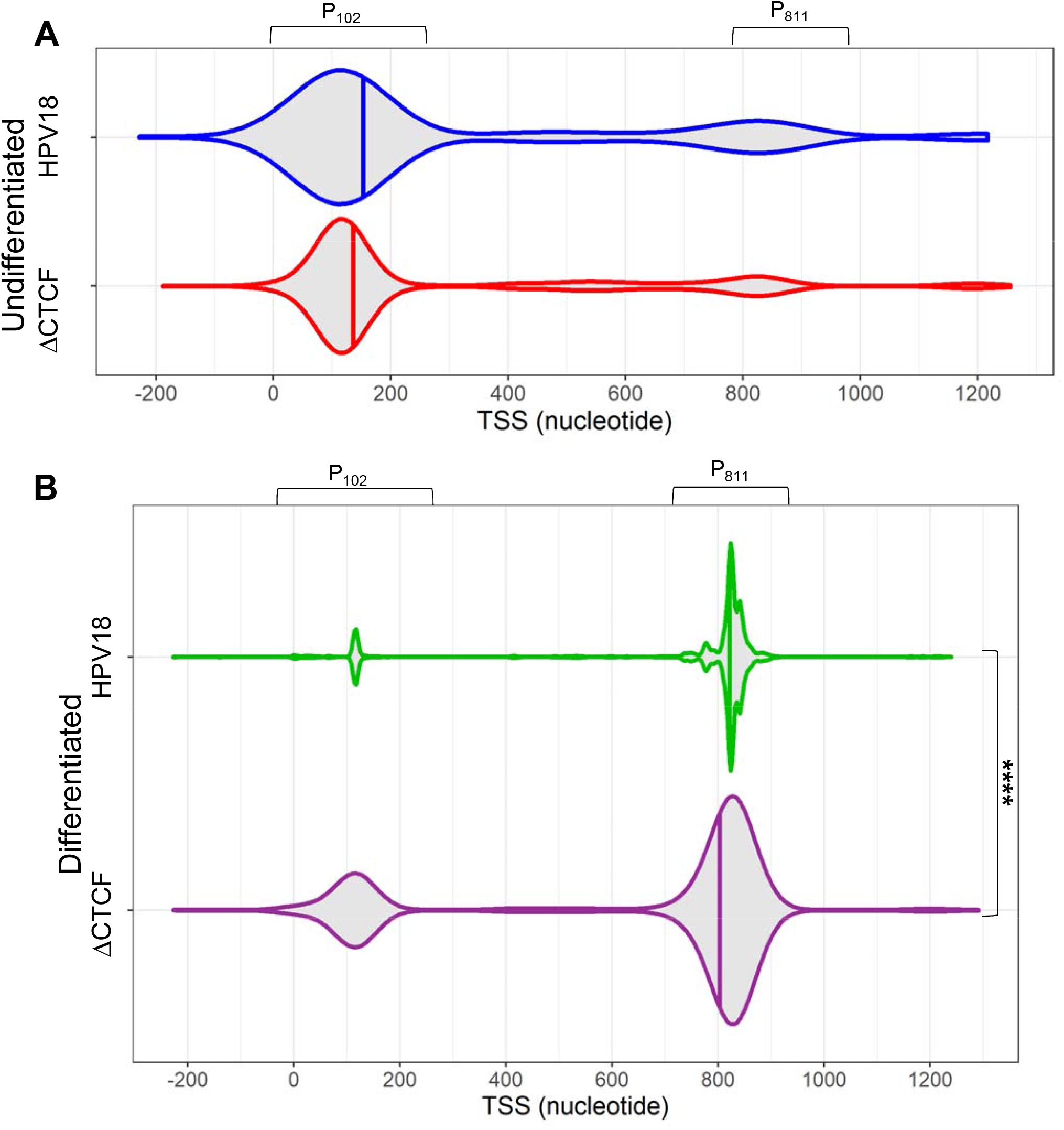
CTCF regulates efficient differentiation-dependent HPV18 late promoter activation. The TSS of each viral transcript was identified and the distribution shown in violin plots in (A) undifferentiated and (B) differentiated keratinocytes containing WT HPV18 (blue and green, respectively) and ΔCTCF-HPV18 (red and purple, respectively) episomes. Data distribution are shown by the kernel shape and median indicated with a vertical solid line. The wider sections of the violin plot indicate a high probability of TSS usage within that region of the HPV18 genome. The shape of the distribution indicates the concentration of data points in a particular region; the steeper the side of each bubble indicates a greater concentration of data points. ns, not significant; ****p<0.0001 (Fisher’s test).

We previously demonstrated that in undifferentiated cells, HPV18-ΔCTCF episomes had increased trimethylation of lysine 4 in histone 3 (H3K4Me3) at the P_102_ early promoter compared to WT, correlating with increased promoter activity. However, while differentiation of HPV18 WT genome-containing cells resulted in a significant enrichment of H3K4Me3 at the P_811_ late promoter, no such enrichment was observed in HPV18-ΔCTCF episomes (6). Enhanced acetylation of histones is also indicative of enhanced activation of transcription by facilitating increased chromatin accessibility and the recruitment of transcriptional activators (41). We therefore assessed the changes in histone 4 acetylation (H4Ac) in HPV18 episomes induced by keratinocyte differentiation. H4Ac abundance in the viral genome in undifferentiated cells was detectable at low levels, consistent with restricted virus transcription (**Fig.6A**). Differentiation of the cells in methylcellulose resulted in a dramatic increase in H4Ac abundance throughout the WT-HPV18 genome, with an over 10-fold enrichment at the P_811_ late promoter, consistent with increased production of late transcripts (**Fig.6A**). However, HPV18-ΔCTCF episomes were devoid of H4Ac with only a small, insignificant increase in H4Ac abundance at the P_811_ following differentiation (**Fig.6B**). Together, these findings suggest that CTCF recruitment to the E2-ORF is necessary for appropriate epigenetic programming of the viral chromatin and differentiation-dependent transcriptional activation of P_811_.

**Figure 6:**
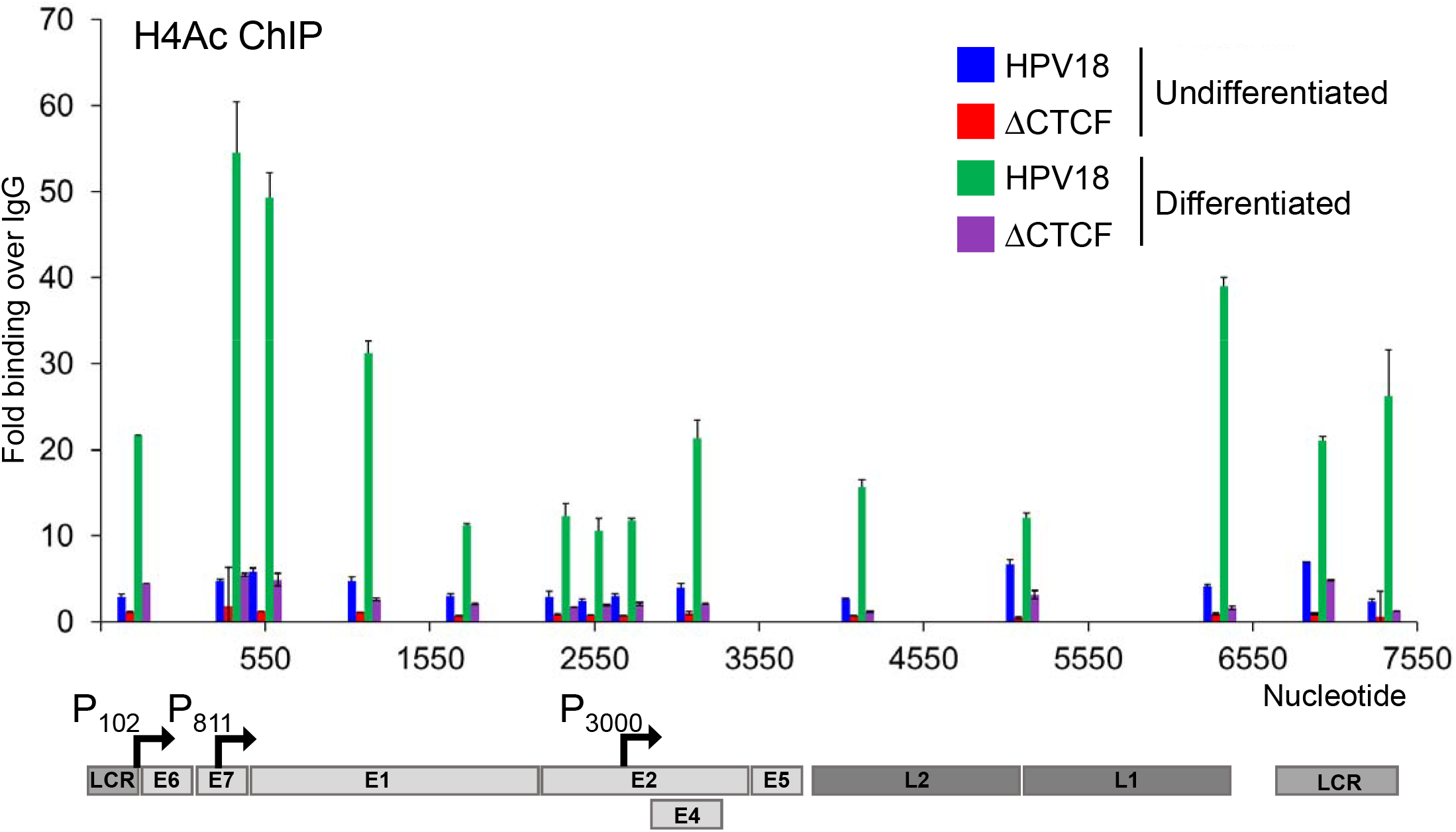
Keratinocyte differentiation induces increased H4Ac abundance at the HPV18 late promoter in wild type but not ΔCTCF-HPV18 genome-containing cells. HPV18 WT and HPV18-ΔCTCF genome-containing primary keratinocytes grown in monolayer (undifferentiated; blue and green, respectively) or differentiated in methylcellulose-containing media for 48 hrs (green and purple, respectively). Enrichment of H4Ac was assessed by ChIP-qPCR. Each bar in the chart represents the mid-point for primer pairs used to amplify immunoprecipitated chromatin. Fold binding over IgG control was calculated. The data shown are the mean and standard deviation of three independent replicates. Annotation of the HPV18 LCR, promoters and ORFs is provided below.

To determine whether the reduced differentiation-dependent activation of P_811_ in HPV18-ΔCTCF genomes resulted in reduced late protein expression, we analysed E1^E4 protein in methylcellulose differentiated cultures. Western blotting of lysates harvested from HPV-18 WT and -ΔCTCF genome containing cells before and after differentiation revealed an induction of involucrin protein expression. However, there was a significant attenuation of E1^E4 protein expression when CTCF binding to the viral genome was abrogated (**Fig.7A and B**). Since L1 protein is not robustly expressed in methylcellulose differentiated keratinocytes, we analysed L1 protein expression by immunostaining organotypic raft culture sections derived from two independent donors of HPV18-WT and -ΔCTCF genome containing cells. L1 positive cells were visible in the upper layers of HPV18-WT genome containing rafts but were barely detectable in HPV18-ΔCTCF rafts and this difference was significant (**Fig.7C and D**). While the total number of E1^E4 positive cells in the upper layers of HPV18-ΔCTCF rafts was not altered, the intensity of E1^E4 staining was notably reduced (**Fig.7C**). Western blot analysis of protein lysates harvested from three independent raft cultures confirmed a significant reduction in E1^E4 protein abundance in HPV18-ΔCTCF genome containing raft cultures in comparison to WT (**Fig.7E**).

**Figure 7:**
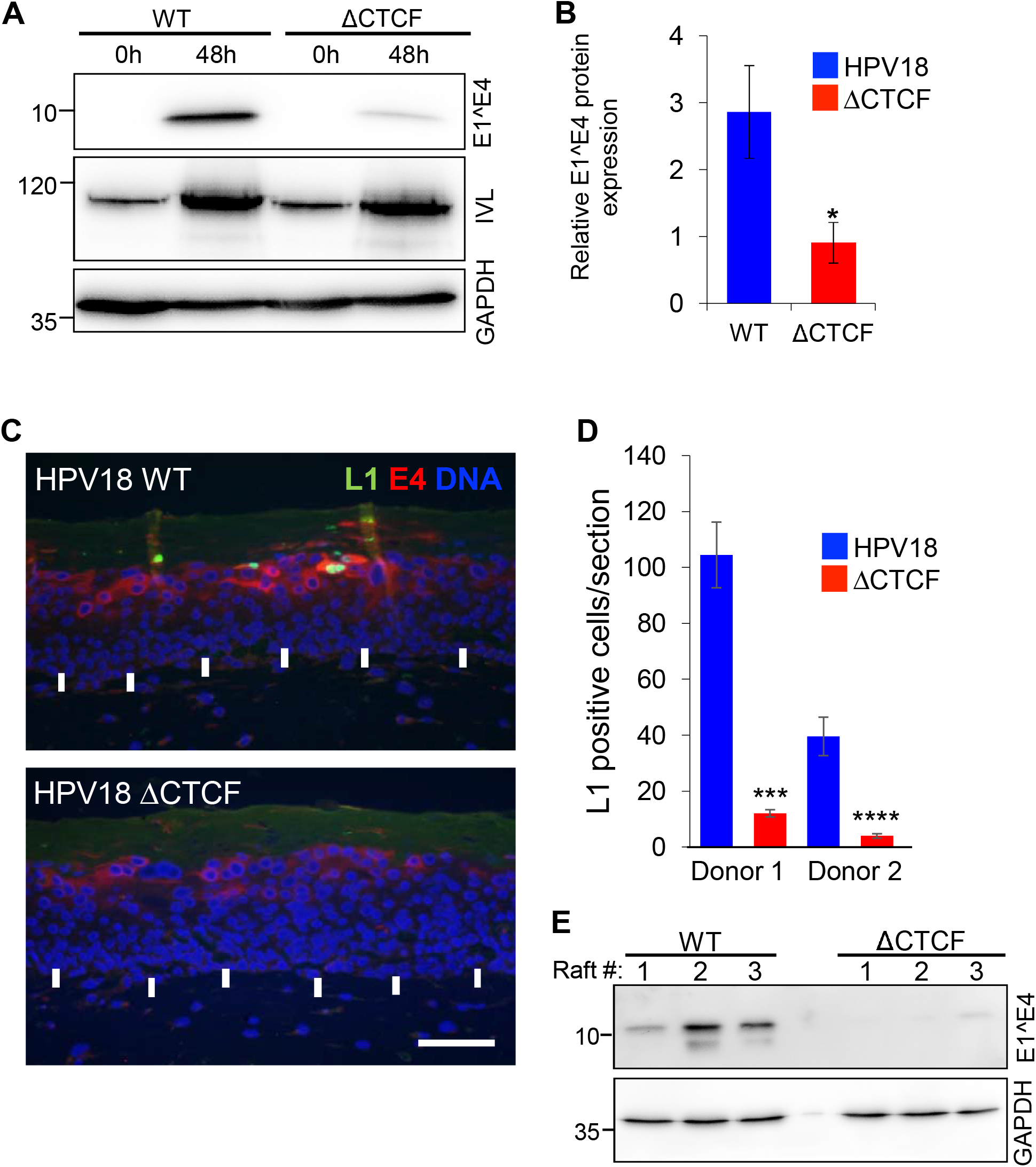
Abrogation of CTCF binding to the HPV18 genome causes a significant reduction in differentiation-dependent late protein abundance. (A) HPV18 genome containing keratinocytes (WT or ΔCTCF) grown in monolayer (undifferentiated, 0h) or differentiated in methylcellulose (48h) and E1^E4, involucrin (IVL) and GAPDH protein expression analysed by Western blotting. Molecular weight markers are indicated on the left (kDa). (B) Relative E1^E4 protein expression in comparison to GAPDH was quantified in three independent experiments by densitometry. Data are the mean +/- standard deviation. * denotes p<0.05. (C) E1^E4 (red) and L1 (green) protein abundance was analysed by indirect immunofluorescence in epithelia derived from wild type and ΔCTCF-HPV18 genome-containing keratinocytes grown in organotypic raft culture. Cellular nuclei are shown in blue, and the basal layer indicated with white arrows. Scale bar indicates 10 μm. (D) The total number of L1 positive cells per section of three independent raft cultures grown from two independent keratinocyte donors was counted. Data show the mean +/- standard deviation. *** p<0.001, **** p<0.0001. (E) E1^E4 protein expression in organotypic raft cultures was assessed by Western blotting lysates harvested from three independent raft cultures alongside GADPH loading control. Molecular weight markers are indicated on the left.

## DISCUSSION

The differentiation-dependent regulation of papillomavirus transcription is fundamental to the productivity and persistence of infection. Previous studies have shown that the viral early (P_105_) promoter is active in basal keratinocytes and becomes further activated as the cells enter terminal differentiation (5, 6). In contrast, the viral late promoter (P_811_) is repressed in undifferentiated basal cells and strongly activated upon induction of cellular differentiation (4, 5, 9, 16, 42). In this study, we have utilised direct, long-read RNA sequencing to quantitatively analyse HPV18 promoter activity and to dissect the role of CTCF in regulating viral transcription at key stages of the virus life cycle. Our findings confirm the differentiation-dependent model of HPV transcription control; transcripts that originate from the P_105_ promoter are dominant in undifferentiated cells and further increased in abundance upon cellular differentiation. The abundance of transcripts originating from the P_811_ late promoter is low in undifferentiated cells but is dramatically upregulated when cells are differentiated. Transcription originating from the P_520_ and P_3000_ promoter regions is also activated by cellular differentiation but overall, these promoters are far less active than either the P_105_ or P_811_ promoters. The P_E8_ promoter is equally weak in both undifferentiated and differentiated cells with only two transcript species that originate from this TSS. The most dominant transcript identified from the P_E8_ promoter was spliced at 1357^3434 and encodes E8^E2 and E5. The second transcript, spliced at 1357^3465 to encode E5 only, was slightly increased in expression in differentiated cell cultures.

Transcripts that encode fusion products between the E2 and E4 ORFs (E2^E4) have been previously described (38). These transcripts were reported to originate upstream of the E2 start code at position 2816 in HPV18 and therefore encode a protein fusion between the N-terminus of E2 and the C-terminus of E4. E2^E4S encoding transcripts, spliced at 2853^3434, were not identified in any of our Nanopore or RNA-Seq datasets. We did however detect transcripts spliced at 3165^3434, which have been previously described to encode a fusion protein termed E2^E4L (38). However, this transcript was detected at very low abundance (~1 RPM) and only in differentiated keratinocytes. Interestingly, we also identified a third transcript that may encode a previously uncharacterised E2^E4 fusion protein, which we termed E2^E4XL. This transcript originated from the P_105_ promoter and was spliced at 929^2779 and 3284^3434. Like E2^E4L, this transcript retains the E2 start codon but potentially encodes amino acids 1-156 of E2 fused to amino acid 6-88 of E4, but it remains to be determined if this transcript encodes a bone fide E2^E4 fusion protein. Interestingly, most of the transcripts that originated from the P_3000_ promoter were also spliced 3165^3434 or 3284^3434. These transcripts were in higher abundance than those originating from the P_105_ promoter in both undifferentiated and differentiated cells, but since they lack the E2 start codon, they are likely to encode E5 protein only. Supporting this hypothesis, splicing of transcripts originating from the P_3000_ promoter 3165^3434 and 3284^3434 respectively removes several intronic ATG codons (7 and 11, respectively), potentially facilitating enhanced translation of E5.

Comparison of the HPV18 transcript map between WT and ΔCTCF genome-containing cells revealed several important phenotypes. Firstly, abrogation of CTCF binding resulted in enhanced production of transcripts originating from the P_105_ promoter, in agreement with our previous findings (6, 7). The increased P_105_ activity resulted in an increase in transcripts spliced at 233^416-929^3434 (encoding E6*I, E7, E1^E4 and E5) and 929^3434, (encoding E6, E7, E1^E4 and E5) while there was a small decrease in transcripts spliced solely at 233^416 (encoding E6*I, E1, E7 and E2) and 233^3434 (the only known transcript to encode E6*II), confirming our previous observation that abrogation of CTCF binding to the HPV18 genome reduces the abundance of transcripts spliced at 233^3434 (7). In addition, a marked decrease in transcripts spliced at 3165^3434 and 3284^3434 was observed in ΔCTCF-HPV18 genome containing cells in comparison to WT, confirming our initial analysis of HPV18 transcript splicing by conventional RNA-Seq. These data indicate that CTCF plays a key role in splice donor choice when splicing to the dominant splice acceptor site at nucleotide 3434 in the HPV18 genome.

A functional role for CTCF in influencing cellular co-transcriptional alternative splicing has been previously demonstrated. CTCF binding within or downstream of weak exons can promote exon inclusion by creating a roadblock to pause RNA polymerase II progression, allowing greater splicing efficiency (22, 23, 43). Interestingly, CTCF-mediated chromatin loop stabilisation between gene promoter and exon regions also plays a key role in regulating alternative splicing events. Exons downstream of a CTCF stabilised promoter-exon loop are more likely to be included in the nascent mRNA, providing a functional link between three-dimensional chromatin organisation and splicing regulation (24). Notably, in this study we show that this mechanism of splicing regulation is recapitulated in the HPV18 genome. CTCF binding within the HPV18 E2 ORF, upstream of weak slice donor sites at 3165 and 3284 results in YY1-dependent stabilisation of a distinct chromatin loop with the upstream viral promoter (6) and correlates with increased splicing at both 3165^3434 and 3284^3434 to produce E5 encoding transcripts.

As expected, cellular differentiation strongly induced P_811_ promoter activation in WT HPV18 episomes. However, HPV18-ΔCTCF genome containing cells displayed a notable reduction in the abundance of transcripts originating from this promoter following differentiation. Differentiation dependent activation of the P_3000_ promoter was also attenuated in virus unable to bind CTCF. In contrast the P_520_ promoter in ΔCTCF-HPV18 was active in both undifferentiated and differentiated cells, albeit at a low level. Activity of P_520_ was induced by cellular differentiation in WT HPV18 cells but was not further activated in ΔCTCF-HPV18 cells. In agreement with the observed differentiation induced activation of the P_811_ and P_3000_ promoters in WT HPV18 episomes, we demonstrated a marked increase in H4Ac enrichment, particularly in around the P_811_ and P_3000_ promoters. Interestingly, H4Ac enrichment following differentiation was not recapitulated in ΔCTCF-HPV18 episomes, indicating that CTCF binding to the E2-ORF is important for enhanced transcriptional activation in the late stages of the virus life cycle. Importantly, attenuation of differentiation-dependent late promoter activation in ΔCTCF-HPV18 resulted in significantly reduced E1^E4 protein expression following methylcellulose differentiation and an almost complete loss of L1 protein expression in stratified epithelia. These results demonstrate for the first time that CTCF has essential functions in differentiation-dependent transcriptional dynamics in the late stages of the HPV life cycle.

## ACKNOWLEDGEMENTS

This work was funded by grants from the Medical Research Council awarded to JLP and SR (MR/R022011/1, MR/T015985/1 and MR/N023498/1). IP was supported by a Cancer Research UK non-clinical PhD studentship awarded to JLP and SR. BN is funded through the Cancer Research UK Birmingham Centre award C17422/A25154. The funders had no role in study design, data collection and interpretation, or the decision to submit the work for publication. We thank Dr. Joseph Spitzer and his patients for the collection and donation of foreskin tissue.

## REFERENCES

1. Tommasino M. 2014. The human papillomavirus family and its role in carcinogenesis. Semin Cancer Biol 26:13–21.

2. Stunkel W, Bernard HU. 1999. The chromatin structure of the long control region of human papillomavirus type 16 represses viral oncoprotein expression. J Virol 73:1918–30.

3. Hummel M, Lim HB, Laimins LA. 1995. Human papillomavirus type 31b late gene expression is regulated through protein kinase C-mediated changes in RNA processing. J Virol 69:3381–8.

4. Ruesch MN, Stubenrauch F, Laimins LA. 1998. Activation of papillomavirus late gene transcription and genome amplification upon differentiation in semisolid medium is coincident with expression of involucrin and transglutaminase but not keratin-10. J Virol 72:5016–24.

5. Wooldridge TR, Laimins LA. 2008. Regulation of human papillomavirus type 31 gene expression during the differentiation-dependent life cycle through histone modifications and transcription factor binding. Virology 374:371–80.

6. Pentland I, Campos-Leon K, Cotic M, Davies KJ, Wood CD, Groves IJ, Burley M, Coleman N, Stockton JD, Noyvert B, Beggs AD, West MJ, Roberts S, Parish JL. 2018. Disruption of CTCF-YY1-dependent looping of the human papillomavirus genome activates differentiation-induced viral oncogene transcription. PLoS Biol 16:e2005752.

7. Paris C, Pentland I, Groves I, Roberts DC, Powis SJ, Coleman N, Roberts S, Parish JL. 2015. CCCTC-binding factor recruitment to the early region of the human papillomavirus 18 genome regulates viral oncogene expression. J Virol 89:4770–85.

8. Beagan JA, Duong MT, Titus KR, Zhou L, Cao Z, Ma J, Lachanski CV, Gillis DR, Phillips-Cremins JE. 2017. YY1 and CTCF orchestrate a 3D chromatin looping switch during early neural lineage commitment. Genome Res doi:10.1101/gr.215160.116.

9. del Mar Pena LM, Laimins LA. 2001. Differentiation-dependent chromatin rearrangement coincides with activation of human papillomavirus type 31 late gene expression. J Virol 75:10005–13.

10. Burley M, Roberts S, Parish JL. 2020. Epigenetic regulation of human papillomavirus transcription in the productive virus life cycle. Semin Immunopathol 42:159–171.

11. Grassmann K, Rapp B, Maschek H, Petry KU, Iftner T. 1996. Identification of a differentiation-inducible promoter in the E7 open reading frame of human papillomavirus type 16 (HPV-16) in raft cultures of a new cell line containing high copy numbers of episomal HPV-16 DNA. J Virol 70:2339–49.

12. Hummel M, Hudson JB, Laimins LA. 1992. Differentiation-induced and constitutive transcription of human papillomavirus type 31b in cell lines containing viral episomes. J Virol 66:6070–80.

13. Wilson R, Fehrmann F, Laimins LA. 2005. Role of the E1--E4 protein in the differentiation-dependent life cycle of human papillomavirus type 31. J Virol 79:6732–40.

14. Peh WL, Brandsma JL, Christensen ND, Cladel NM, Wu X, Doorbar J. 2004. The viral E4 protein is required for the completion of the cottontail rabbit papillomavirus productive cycle in vivo. J Virol 78:2142–51.

15. Bodily JM, Meyers C. 2005. Genetic analysis of the human papillomavirus type 31 differentiation-dependent late promoter. J Virol 79:3309–21.

16. Songock WK, Scott ML, Bodily JM. 2017. Regulation of the human papillomavirus type 16 late promoter by transcriptional elongation. Virology 507:179–191.

17. Wang X, Meyers C, Wang HK, Chow LT, Zheng ZM. 2011. Construction of a full transcription map of human papillomavirus type 18 during productive viral infection. J Virol 85:8080–92.

18. Toots M, Mannik A, Kivi G, Ustav M, Jr., Ustav E, Ustav M. 2014. The transcription map of human papillomavirus type 18 during genome replication in U2OS cells. PLoS One 9:e116151.

19. Mole S, McFarlane M, Chuen-Im T, Milligan SG, Millan D, Graham SV. 2009. RNA splicing factors regulated by HPV16 during cervical tumour progression. J Pathol 219:383–91.

20. McFarlane M, MacDonald AI, Stevenson A, Graham SV. 2015. Human Papillomavirus 16 Oncoprotein Expression Is Controlled by the Cellular Splicing Factor SRSF2 (SC35). J Virol 89:5276–87.

21. Shukla S, Kavak E, Gregory M, Imashimizu M, Shutinoski B, Kashlev M, Oberdoerffer P, Sandberg R, Oberdoerffer S. 2011. CTCF-promoted RNA polymerase II pausing links DNA methylation to splicing. Nature 479:74–79.

22. Lopez Soto EJ, Lipscombe D. 2020. Cell-specific exon methylation and CTCF binding in neurons regulate calcium ion channel splicing and function. Elife 9.

23. Agirre E, Bellora N, Allo M, Pages A, Bertucci P, Kornblihtt AR, Eyras E. 2015. A chromatin code for alternative splicing involving a putative association between CTCF and HP1alpha proteins. BMC Biol 13:31.

24. Ruiz-Velasco M, Kumar M, Lai MC, Bhat P, Solis-Pinson AB, Reyes A, Kleinsorg S, Noh KM, Gibson TJ, Zaugg JB. 2017. CTCF-Mediated Chromatin Loops between Promoter and Gene Body Regulate Alternative Splicing across Individuals. Cell Syst 5:628–637 e6.

25. Goodwin S, McPherson JD, McCombie WR. 2016. Coming of age: ten years of next-generation sequencing technologies. Nature Reviews Genetics 17:333–351.

26. van Dijk EL, Jaszczyszyn Y, Naquin D, Thermes C. 2018. The Third Revolution in Sequencing Technology. Trends in Genetics 34:666–681.

27. Wilson R, Laimins LA. 2005. Differentiation of HPV-containing cells using organotypic “raft” culture or methylcellulose. Methods Mol Med 119:157–69.

28. Roberts S, Hillman ML, Knight GL, Gallimore PH. 2003. The ND10 Component Promyelocytic Leukemia Protein Relocates to Human Papillomavirus Type 1 E4 Intranuclear Inclusion Bodies in Cultured Keratinocytes and in Warts. Journal of Virology 77:673–684.

29. Günther T, Fröhlich J, Herrde C, Ohno S, Burkhardt L, Adler H, Grundhoff A. 2019. A comparative epigenome analysis of gammaherpesviruses suggests cis-acting sequence features as critical mediators of rapid polycomb recruitment. PLOS Pathogens 15:e1007838.

30. Langmead B, Trapnell C, Pop M, Salzberg SL. 2009. Ultrafast and memory-efficient alignment of short DNA sequences to the human genome. Genome Biol 10:R25.

31. Günther T, Grundhoff A. 2010. The epigenetic landscape of latent Kaposi sarcoma-associated herpesvirus genomes. PLoS Pathog 6:e1000935.

32. Dobin A, Davis CA, Schlesinger F, Drenkow J, Zaleski C, Jha S, Batut P, Chaisson M, Gingeras TR. 2012. STAR: ultrafast universal RNA-seq aligner. Bioinformatics 29:15–21.

33. Thompson S, Thompson S, Cazier J. 2019. CaStLeS (Compute and Storage for the Life Sciences): a collection of compute and storage resources for supporting research at the University of Birmingham. Zenodo.

34. Schwenzer H, Abdel Mouti M, Neubert P, Morris J, Stockton J, Bonham S, Fellermeyer M, Chettle J, Fischer R, Beggs AD, Blagden SP. 2021. LARP1 isoform expression in human cancer cell lines. RNA Biol 18:237–247.

35. Li H. 2018. Minimap2: pairwise alignment for nucleotide sequences. Bioinformatics 34:3094–3100.

36. Reynolds G, Deshmukh NS, Mangham D. 2000. Agitated low temperature epitope retrieval (ALTER): Effective antigen retrieval for immunohistochemistry with excellent morphological preservation. The Journal of Pathology 190:51A–51A.

37. Mehta K, Gunasekharan V, Satsuka A, Laimins LA. 2015. Human papillomaviruses activate and recruit SMC1 cohesin proteins for the differentiation-dependent life cycle through association with CTCF insulators. PLoS Pathog 11:e1004763.

38. Tan CL, Gunaratne J, Lai D, Carthagena L, Wang Q, Xue YZ, Quek LS, Doorbar J, Bachelerie F, Thierry F, Bellanger S. 2012. HPV-18 E2^E4 chimera: 2 new spliced transcripts and proteins induced by keratinocyte differentiation. Virology 429:47–56.

39. Smits P, Poumay Y, Karperien M, Tylzanowski P, Wauters J, Huylebroeck D, Ponec M, Merregaert J. 2000. Differentiation-dependent alternative splicing and expression of the extracellular matrix protein 1 gene in human keratinocytes. J Invest Dermatol 114:718–24.

40. Donovan-Banfield Ia, Turnell AS, Hiscox JA, Leppard KN, Matthews DA. 2020. Deep splicing plasticity of the human adenovirus type 5 transcriptome drives virus evolution. Communications Biology 3:124.

41. LeRoy G, Rickards B, Flint SJ. 2008. The double bromodomain proteins Brd2 and Brd3 couple histone acetylation to transcription. Mol Cell 30:51–60.

42. Spink KM, Laimins LA. 2005. Induction of the human papillomavirus type 31 late promoter requires differentiation but not DNA amplification. J Virol 79:4918–26.

43. Shukla S, Kavak E, Gregory M, Imashimizu M, Shutinoski B, Kashlev M, Oberdoerffer P, Sandberg R, Oberdoerffer S. 2011. CTCF-promoted RNA polymerase II pausing links DNA methylation to splicing. Nature 479:74–9.

